# High throughput screen identifies IFN-γ-dependent inhibitors of *Toxoplasma gondii* growth

**DOI:** 10.1101/336818

**Authors:** Joshua B. Radke, Kimberly L. Carey, Subrata Shaw, Shailesh R. Metkar, Carol Mulrooney, Jennifer P. Gale, Joshua A. Bittker, Robert Hilgraf, Eamon Comer, Stuart L. Schreiber, Herbert W. Virgin, Jose R. Perez, L. David Sibley

## Abstract

*Toxoplasma gondii* is an obligate intracellular parasite capable of causing severe disease due to congenital infection and in patients with compromised immune systems. Control of infection is dependent on a robust Th1 type immune response including production of interferon gamma (IFN-γ), which is essential for control. IFN-γ activates a variety of anti-microbial mechanisms in host cells, which are then able to control intracellular parasites such as *T. gondii*. Despite the effectiveness of these pathways in controlling acute infection, the immune system is unable to eradicate chronic infections that can persist for life. Similarly, while antibiotic treatment can control acute infection, it is unable to eliminate chronic infection. To identify compounds that would act synergistically with IFN-γ, we performed a high-throughput screen of diverse small molecule libraries to identify inhibitors of *T. gondii*. We identified a number of compounds that inhibited parasite growth *in vitro* at low μM concentrations and that demonstrated enhanced potency in the presence of low level of IFN-γ. A subset of these compounds act by enhancing the recruitment of LC3 to the parasite-containing vacuole, suggesting they work by an autophagy-related process, while others were independent of this pathway. The pattern of IFN-γ-dependence was shared among the majority of analogs from 6 priority scaffolds and analysis of structure activity relationships for one such class revealed specific stereochemistry associated with this feature. Identification of these IFN-γ-dependent leads may lead to development of improved therapeutics due to their synergistic interactions with immune responses.

## Introduction

*Toxoplasma gondii* is a widespread protozoal infection of animals that is transmitted by cats, which serve as the definitive host and shed oocysts into the environment (1). Wild and domesticated herbivorous animals are readily infected by ingestion of oocysts, which are extremely resistant to environmental conditions (1). Following a brief acute phase during which the parasite propagates as fast growing tachyzoites that disseminate to all organs of the body, the parasite differentiates into slow growing bradyzoites that reside in long-lived tissue cysts which form in differentiated cells such as neurons and muscle cells (2). Infections can be passed vertically (3), or through oral ingestion of tissue cysts due to omnivorous or carnivorous feeding (4). Humans often become infected due to water or food born contamination (5, 6) or due to congenital infection (7). Although most infection are benign, a large proportion of the world’s human population is chronically infected (8), putting them at risk of reactivation in the event of a decline in immune surveillance (9)

Interferon gamma (IFN-γ) is the major cytokine responsible for driving the innate immune response to many intracellular pathogens (10), including *Toxoplasma gondii* (11, 12). There are multiple, overlapping pathways attributed to IFN-γ-dependent control of *T. gondii* growth in human cells, and the extent to which these operate in different cell types varies (13). Pathways for IFN-γ-dependent control include induction of a non-canonical autophagy pathway that limits growth (14), starvation by tryptophan degradation in the host cell (15), upregulation of guanylate binding proteins that are recruited to the parasite containing vacuole (16), ubiquitination and lysosomal clearance (17), and induction of premature parasite egress (18). Although the strong Th1 immune response associated with control of toxoplasmosis is beneficial, it also results in immunopathology due to inflammation, a syndrome commonly seen in ocular toxoplasmosis (19). Current therapies for treatment of toxoplasmosis rely on inhibition of the folate pathway in the parasite, although macrolide antibiotics have also been used with some success (20). Although these treatments are designed to block DNA replication and protein synthesis in the parasite, respectively, they are not effective in eliminating the tissue cyst forms that are responsible for chronic infection (21). The effectiveness of these treatments is augmented by the immune response (22), which contributes to control of acute infection but also cannot eradicate the chronic infection (23).

To take advantage of the potential synergy between innate immunity and chemotherapy, we designed a screen to evaluate compounds for their ability to enhance the IFN-γ-dependent host response to *T. gondii* infection. Such combination treatments could potentially allow control at lower drug exposure and with fewer side effects associated with strong Th1 responses. The Broad Institute has previously synthesized close to 100,000 complex small molecules through diversity-oriented synthesis (DOS) for use in high throughput screening (HTS) (24, 25). This DOS collection consists of a variety of different chemical skeletons and they are generally rich in sp3-hybridized carbon atoms (for increased ‘natural product-likeness’) compared to conventional small molecule libraries. We took advantage of this DOS collection and also screened inhibitors from commercial vendors to identify several chemical scaffolds that show enhanced IFN-γ-dependent control of *T. gondii* replication.

## Results and Discussion

### Development of a HTS for IFN-γ-dependent inhibitors of *T. gondii* growth

In order to identify compounds that would enhance the ability of IFN-γ to control *Toxoplasma* infection, two critical components were needed. First, we needed to identify the concentration of IFN-γ that would partially activate cells but not result in complete growth inhibition. Although recent studies indicate various concentrations restrict *T.gondii* growth in human cells (14, 18), the concentration of IFN-γ required to partially restrict growth has not been identified. We determined the concentration of recombinant human IFN-γ required to limit replication of *T. gondii* in HeLa cells that were pre-activated for 24 h with different doses of human IFN-γ. The dose response curve exhibited a very sharp transition from low inhibition at low IFN-γ concentrations that very rapidly increased to maximum inhibition (Fig. 1A). Based on this response, we designed the primary screen and follow up validation assays to use doses of IFN-γ that resulted in an EC_10-20_ (Fig. 1A). Second, to facilitate screening of larger compound collections, we miniaturized a growth assay based on a luciferase expressing parasite line to fit a 384-well format congruent with high throughput screen (HTS) platforms (Fig. 1B). We also added a 4h pre-incubation treatment to ensure compounds had the opportunity to promote cellular responses (Fig. 1B) prior to addition of *T. gondii* parasites. We then used this HTS to evaluate ∼73,000 compounds to identify those that would enhance the innate immune response to *T. gondii* infection. First, all ∼73,000 compounds were screened in duplicate for growth inhibition (single point, 10 μM) in the presence of a low dose of IFN-γ (1.25 U/mL). To ensure the compounds were targeting the parasite and not the host cell, a cytotoxicity screen against HeLa cells was conducted in parallel (Fig. 2). Removal of the cytotoxic compounds left 2,143 compounds that inhibited *T. gondii* growth by >50% at 10 μM, and < 50% host cell growth inhibition at a concentration of 20 μM. Next, these compounds were retested using an eight-point dose response growth inhibition assay to identify 880 active compounds that were then tested in the absence of IFN-γ (Fig. 2). Comparison of these activities identified 80 compounds with an EC_50_ values for parasite growth inhibition of < 10 μM, > 2 fold IFN-γ dependence, and toxicity to host cells > 20 μM (Fig. 2). Structural clustering and structurally similar analogs were used to further refine the hit list to 72 compounds that showed IFN-γ-dependence (Fig. 2, see Table S1 for complete data). A summary of the screening tree and compound prioritization is defined in Figure 2.

**Figure 1:**
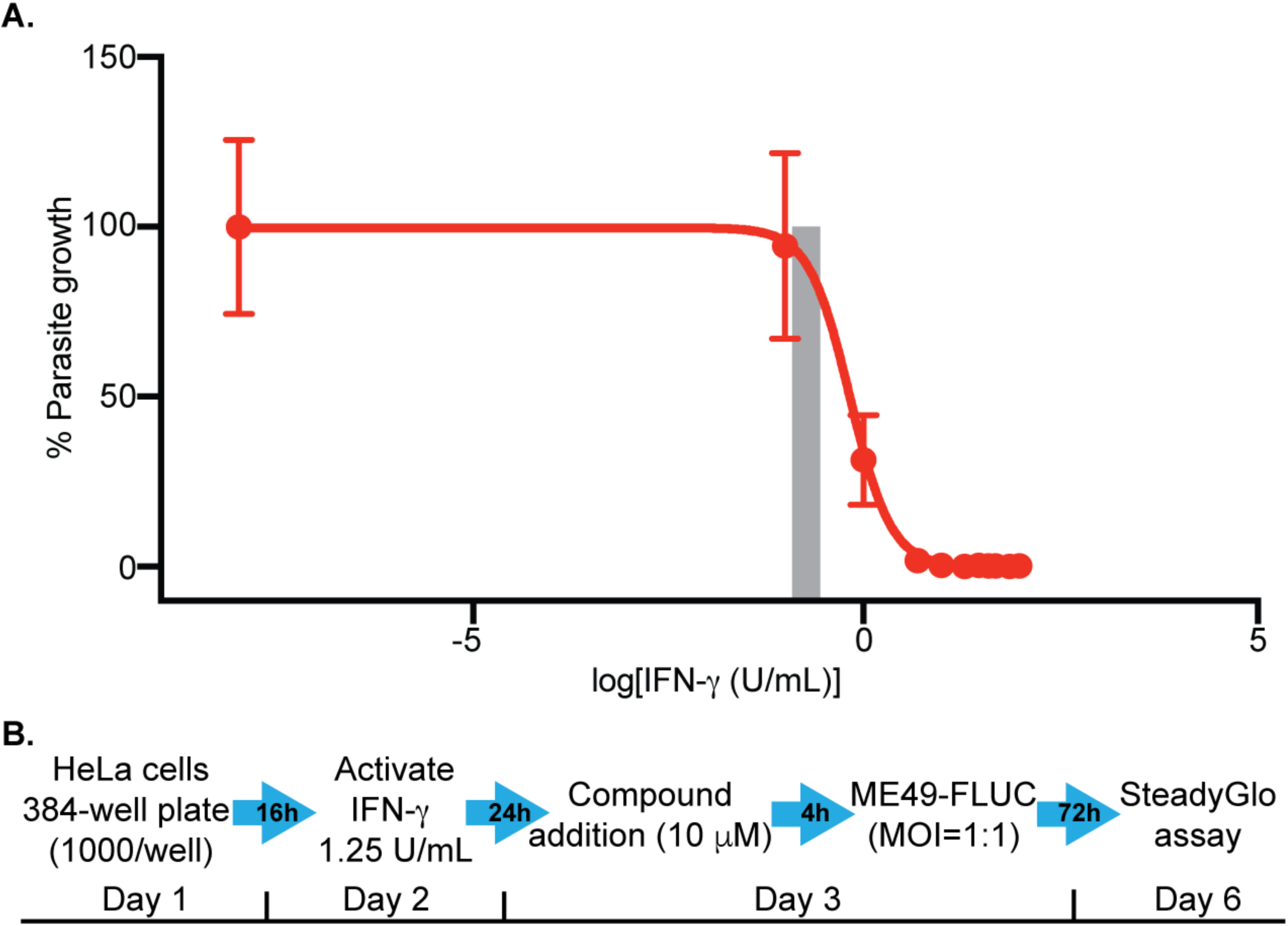
Titration of the effects of IFN-γ on *T. gondii* growth. A dose response curve for inhibition of *T. gondii* tachyzoite growth in response to increasing concentration of IFN-γ. HeLa cells were activated for 24 h with recombinant human IFN-γ (from 100 U/mL to 0.1 U/mL) prior to infection with tachyzoites of the ME49-FLUC strain. Growth was measured by luciferase activity at 72 h post-infection. Data presented as percent relative light units (% RLU) normalized to growth in naïve (unactivated) HeLa cells. Three biological replicates each with technical replicates, N = 9. Shaded region corresponds to the EC_10_-EC_20_. **B)** Diagram of HTS to monitor inhibition of *T. gondii* growth in the presence of low dose IFN-γ. HeLa cells were plated in 384-well plates, allowed to settle and activated with low dose IFN-γ (1.25 U/mL) for 24 h prior to addition of compounds (10 μM). Compounds were allowed 4 h incubation prior to the addition of ME49-FLUC strain parasites. Growth was measured 72 h post infection based on the SteadyGlo luciferase assay.

**Figure 2:**
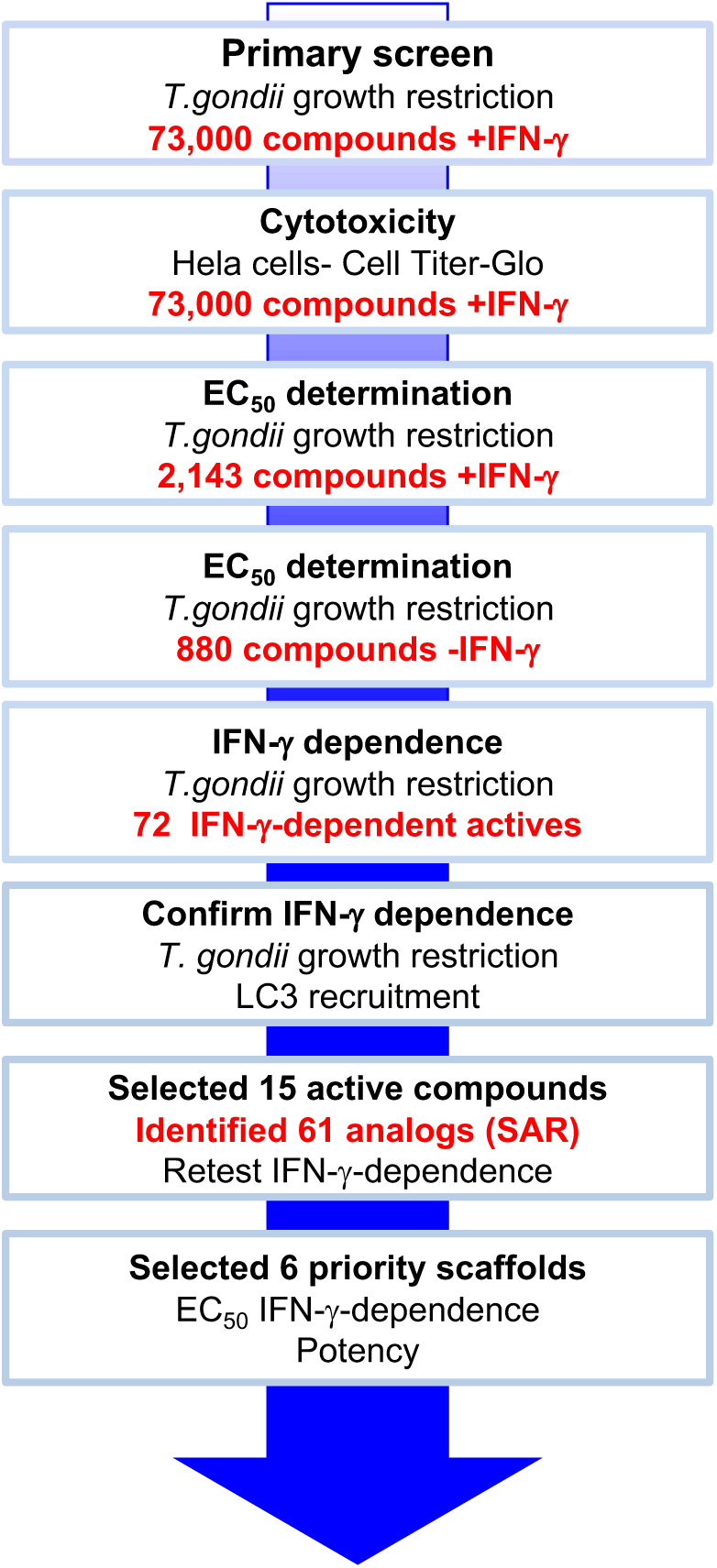
Overview of screen for compound identification. The primary screen evaluated 73,000 molecules for their capacity to restrict *T. gondii* tachyzoite replication *in vitro* (10 μM) using a luciferase-based growth assay. In parallel, the library was screened for cytotoxicity against HeLa cells (20 μM) to remove molecules that were toxic. Following the primary screen, 2,143 molecules were evaluated in an eight-point dose response assay in the presence of IFN-γ (1.25 U/mL) that identified 880 compounds with EC_50_ ≤ 10 μM. These 880 active compounds were retested in an eight-point dose response in the absence of IFN-γ. A total of 72 compounds that displayed IFN-γ-dependent growth inhibition were selected for further validation. After retesting, 15 compounds were selected for further study based on IFN-γ-dependent *T. gondii* growth restriction and enhanced recruitment of GFP-LC3 to the parasite containing vacuole. These core scaffolds were used to identify 61 analogs based on structure-activity relationship (SAR). Finally, 6 priority scaffolds were selected for further interrogation of IFN-γ-dependent potency against *T. gondii.*

### Downstream assays to validate *T. gondii* inhibitors identified in the primary screen

In order to further evaluate the 72 compounds identified in the primary screen, compounds were re-supplied to assure batch to batch consistency and tested in a single point assay (5 μM compound) in the presence vs. absence of low dose IFN-γ using the schematic shown in Figure 3A (Table S1). In total, 35 of 72 compounds were validated to inhibit growth of the tachyzoite stage of *T. gondii* in an IFN-γ-dependent manner, although they differed in the fold change in the presence vs. absence of IFN-γ (Table S1). Vacuoles containing *T. gondii* parasites are normally devoid of LC3, but recruitment is dramatically enhanced by treatment with IFN-γ, resulting in growth impairment (14). Hence, one potential mechanism for the IFN-γ-dependent inhibition would be increased recruitment of LC3. To determine the effect of the 72 compounds on LC3 recruitment, we developed an image-based screen to evaluate recruitment of GFP-LC3 to the parasite-containing vacuole (Fig. 3B). Quantification of GFP-LC3 positive vacuoles revealed that in the presence of high levels of IFN-γ (50 U/mL), 12.4% of parasite-containing vacuoles were GFP-LC3 positive, compared with 0.63% of positive parasite-containing vacuoles in a naïve HeLa cells (Fig. 3C). When evaluated using this image-based assay, a number of compounds enhanced LC3 recruitment to parasite containing vacuole (Fig. 4). These inhibitors were grouped into three sets, depending on the extent to which they were IFN-γ dependent, enhanced LC3, or were simply growth inhibitors (Fig 4A), as described further below. Enhancement of LC3 recruitment was positively correlated with growth inhibition (Fig. 4B). This pattern suggests that these compounds may act by enhancing the recruitment of a non-canonical autophagy pathway that was previously described to inhibit *T. gondii* growth in HeLa cells (14, 17). However, among the most active inhibitors, there was not a strong correlation between LC3 recruitment and IFN-γ-dependent inhibition (Fig. 4C). Taken together, these findings indicate that the inhibitory properties of these compounds can be attributed both to enhanced LC3 recruitment and to IFN-γ dependence, yet these two pathways are largely independent.

**Figure 3:**
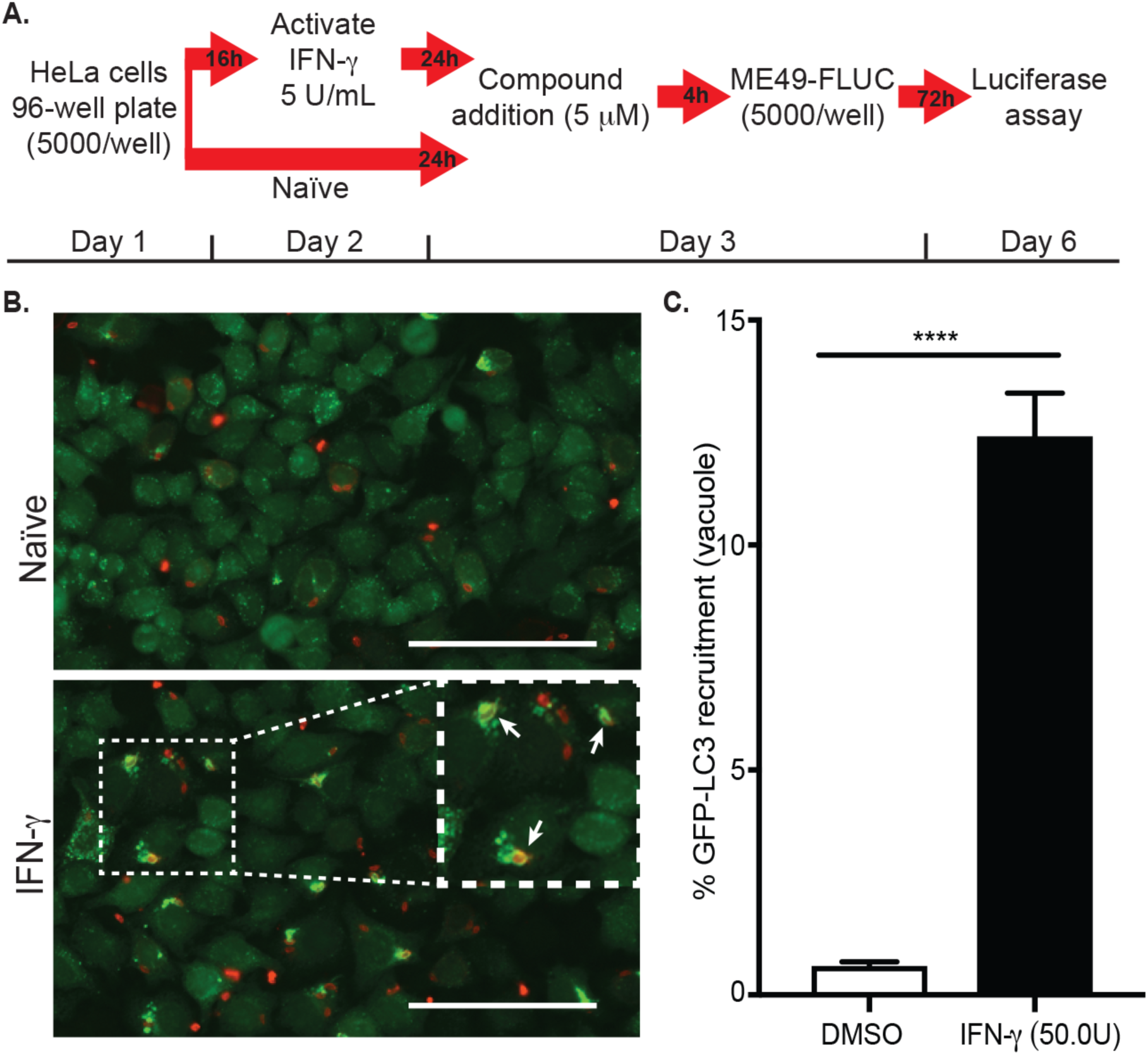
Confirmation of IFN-γ dependence and recruitment of GFP-LC3. **A)** Diagram of single-point inhibition assay for confirming IFN-γ dependence. Hela cells were plated and settled for 16 h, then either left naïve or activated for 24 h with 5 U/mL IFN-γ prior to addition of compounds (5 μM) for 4 h. ME49-FLUC parasites (5,000 / well) were added and luciferase activity was determined at 72 h post-infection. Compounds were screened in duplicate and the average percent inhibition compared to naïve or 5 U/mL IFN-γ activated control cells (DMSO vehicle only) was determined. **B)** Localization of GFP-LC3 in naïve or IFN-γ activated HeLa cells 6h post-infection with ME49-FLUC parasites. GFP-LC3 was localized with mouse anti-GFP antibody (green) and the parasite was identified with polyclonal rabbit anti-RH antibody (red). Images were collected using a Cytation 3 Imaging Multi-Mode Imager, 20x magnification. Inset indicates co-localization of GFP-LC3 (green) with *T. gondii* containing vacuoles (red) (white arrows). Scale bars = 100 microns. **C)** Quantitative fluorescence imaging of GFP-LC3 recruitment to the *T. gondii* PVM in naïve or IFN-γ activated (50 U/mL) GFP-LC3 HeLa cells. Percent recruitment was calculated as GFP positive vacuoles / total vacuoles x 100 in each indicated condition. Values shown are mean ± SEM of 3 biological replicates (*P*< 0.0001, Unpaired Student’s t-test), with a minimum of 500 total vacuoles/replicate.

**Figure 4:**
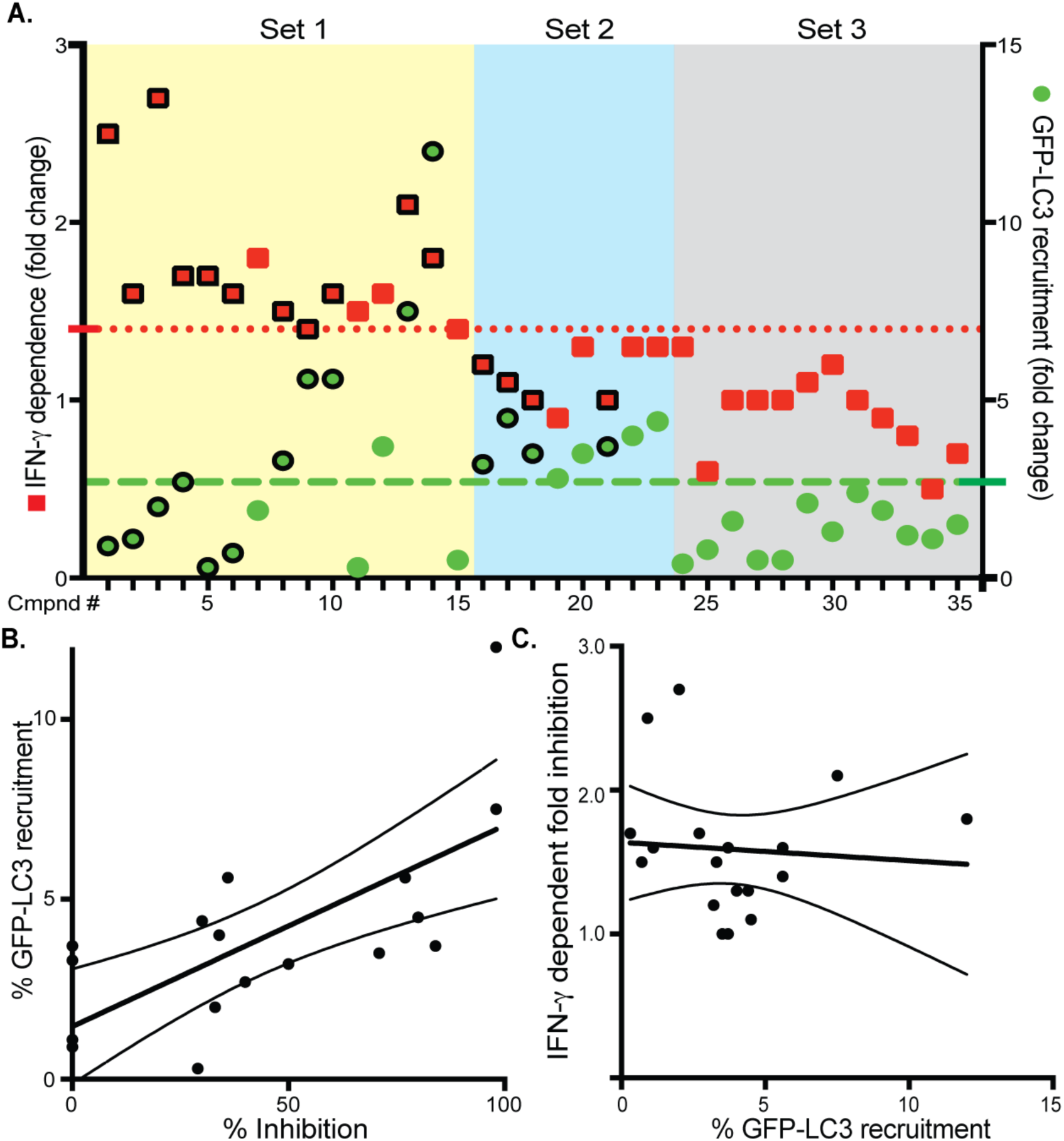
Classification of *T. gondii* inhibitors from IFN-γ dependent growth screen. **A**) IFN-γ-dependent compounds were rescreened at 5 μM and 5 U/ml IFN-γ for growth inhibition and GFP-LC3 recruitment. Compounds were classified based on fold change inhibition ± IFN-γ (fold change ≥ 1.4) or enhanced GFP-LC3 recruitment (fold change ≥ 2.7) to the parasite containing vacuole resulting in three sets as illustrated: Set 1 (15 compounds, yellow box), Enhanced IFN-γ-dependent growth inhibition; Set 2 (8, blue box), Increased GFP-LC3 recruitment to the PVM; Set 3 (12, gray box), Inhibition of *T. gondii* growth independent of IFN-γ or GFP-LC3 recruitment. Data shown are averages of three biological replicates. **B**) Linear regression (r^2^ = 0.503) analysis of GFP-LC3 recruitment vs. % inhibition for compounds shown in A. Solid line shows linear regression with 95% confidence interval shown in dashed lines. **C**) Linear regression (r^2^ = 0.006) analysis for IFN-γ dependent fold-growth inhibition vs. % GFP-LC3 recruitment in the absence of INF-γ for compounds shown in A. Solid line shows linear regression with 95% confidence interval shown in dashed lines. Compounds with symbols shown in bold were selected for further analysis.

The 35 compounds that were validated in these secondary screens were grouped into three sets based on their inhibition of growth, dependence on IFN-γ and involvement of LC3 recruitment. Set one contains 15 compounds that show IFN-γ-dependent inhibition of 1.4 fold or greater (Fig. 4A, Table S1), and approximately half these show enhanced LC3 recruitment (Fig. 4A). Set two contains 8 compounds that show enhanced LC3 recruitment but do not show marked IFN-γ-dependence (Fig. 4A). Finally, set 3 contains 12 potent inhibitors that are neither IFN-γ-dependent nor enhance LC3 (Fig. 4A). Because our primary interest was to identify compounds that were IFN-γ-dependent, we selected 15 compounds from sets 1 (n=11) and 2 (n=4) for further study (identified in black outlined symbols in Fig. 4A, and listed in Table S1) based on these properties as well as the chemical scaffolds they represent.

### SAR evaluation of lead compound series from IFN-γ dependent growth of inhibitors of *T. gondii*

To better understand the structural features important for IFN-γ-dependent growth inhibition of *T. gondii*, 61 analogs centered on the 15 prioritized hits were selected based on stereochemical properties and structural diversity (Table S2). We retested all 76 compounds (15 initial hits and 61 analogs) in a single point screen using 10 μM compound in the presence and absence of low dose IFN-γ (5 U/mL) in order to identify compound series that illustrated IFN-γ dependence across the series (Fig. 5, Table S2). We visualized IFN-γ dependence for each molecule by plotting inhibition in the presence vs. absence of IFN-γ (Fig. 5). These values were compared to a regression line defined by a growth-inhibiting dose of pyrimethamine (3 μM) and normal growth in a naïve HeLa cells (black line, Fig. 5). Compounds that fall below the regression line showed a modest level of IFN-γ-dependence (red arrow) and the most potent molecules fall nearest the origin (black arrow). Interestingly, the majority of compounds from 6 chemical series fell outside the 95% confidence interval (CI, dashed black line) of the regression line as highlighted by the red quadrilateral in Fig. 5. These 6 compound series that together comprise 32 analogs were selected for further evaluation in the presence and absence of low dose IFN-γ (Table S3). Testing EC_50_ values confirmed that 5 of the 6 series displayed IFN-γ dependence including those series defined by the parent compounds BRD1354, BRD1871, BRD5675, BRD7522, and BRD0036 (Table S3). Of these compounds, three of them (i.e. BRD1354, BRD1871 and BRD5675) were also found to enhance GFP-LC3 recruitment (cutoff ≥ 2.7 fold, Table S1). These findings suggest that the activities of these compounds may involve other ATG proteins, although further studies would be required to confirm this dependence. Two chemical series, BRD1871 and BRD7522, were selected to further illustrate SAR patterns that govern IFN-γ dependence in enhancing *T. gondii* growth inhibition.

**Figure 5:**
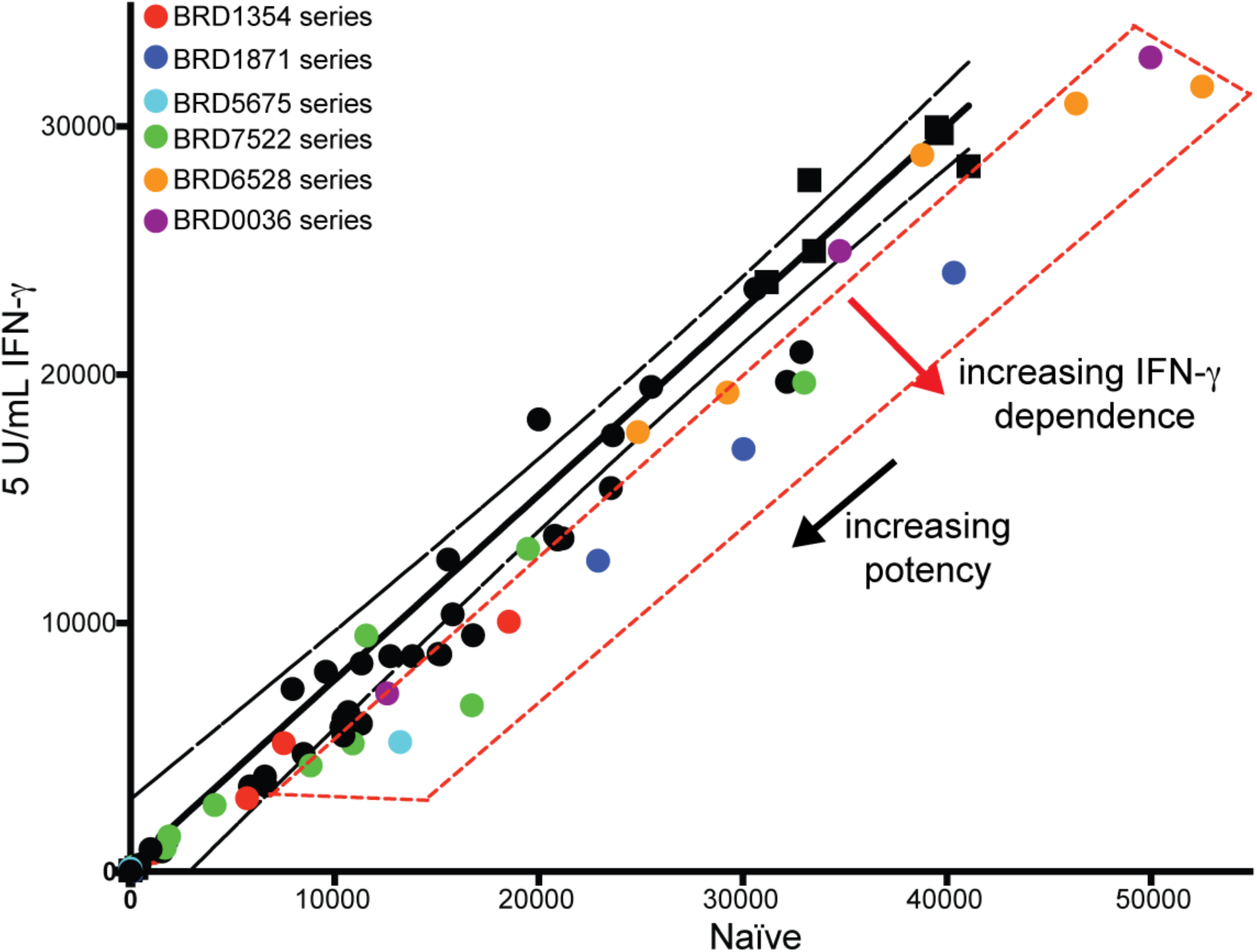
Comparison of compound series for IFN-γ-dependent growth inhibition of *T. gondii*. A total of 61 total analogs together with 15 parent compounds from Sets 1 and 2 above were retested for IFN-γ-dependent growth of *T. gondii* in a replicate screen (10 μM compound, ± 5 U/mL IFN-γ). Plot of average replicate values for naive vs. low dose IFN-γ activated cells for all 76 compounds. The solid line is a linear regression of growth inhibition with 3 μM pyrimethamine (positive control) vs. DMSO (negative control), where dotted lines indicate the 95% confidence interval. Compounds that fall within the red box demonstrated IFN-γ-dependent growth inhibition. Complete data for all 76 compounds can be found in Table S2.

### Compound series BRD1871 has three analogs that exhibit IFN-γ-dependent growth inhibition

The bridged bicyclic azetidine compound series exemplified by one of the original hits compound BRD1871 contained three stereochemical analogs that illustrated increased potency against *T. gondii* in the presence of low dose IFN-γ (blue bars) than in naïve HeLa cells (green bars) (Fig. 6A). Due to its potency, we could not determine IFN-γ dependence of BRD4137 in a single point assay, so we determined EC_50_ values to determine the level of IFN-γ-dependence of each compound (Fig. 6C). Both the parent compound and BRD4137 inhibited growth in naïve HeLa cells (BRD1871, EC_50_=10.16 μM, solid red line; BRD4137, EC_50_= 2.21 μM, solid blue line) and were more potent in the presence of IFN-γ (BRD1871, EC_50_ =7.60 μM, dashed red line; BRD4137, EC_50_ = 1.10 μM, dashed blue line). The primary change in the EC_50_ is driven by a change in the Hill slope, which might be explained by heterogeneity in cellular responses (26).

**Figure 6:**
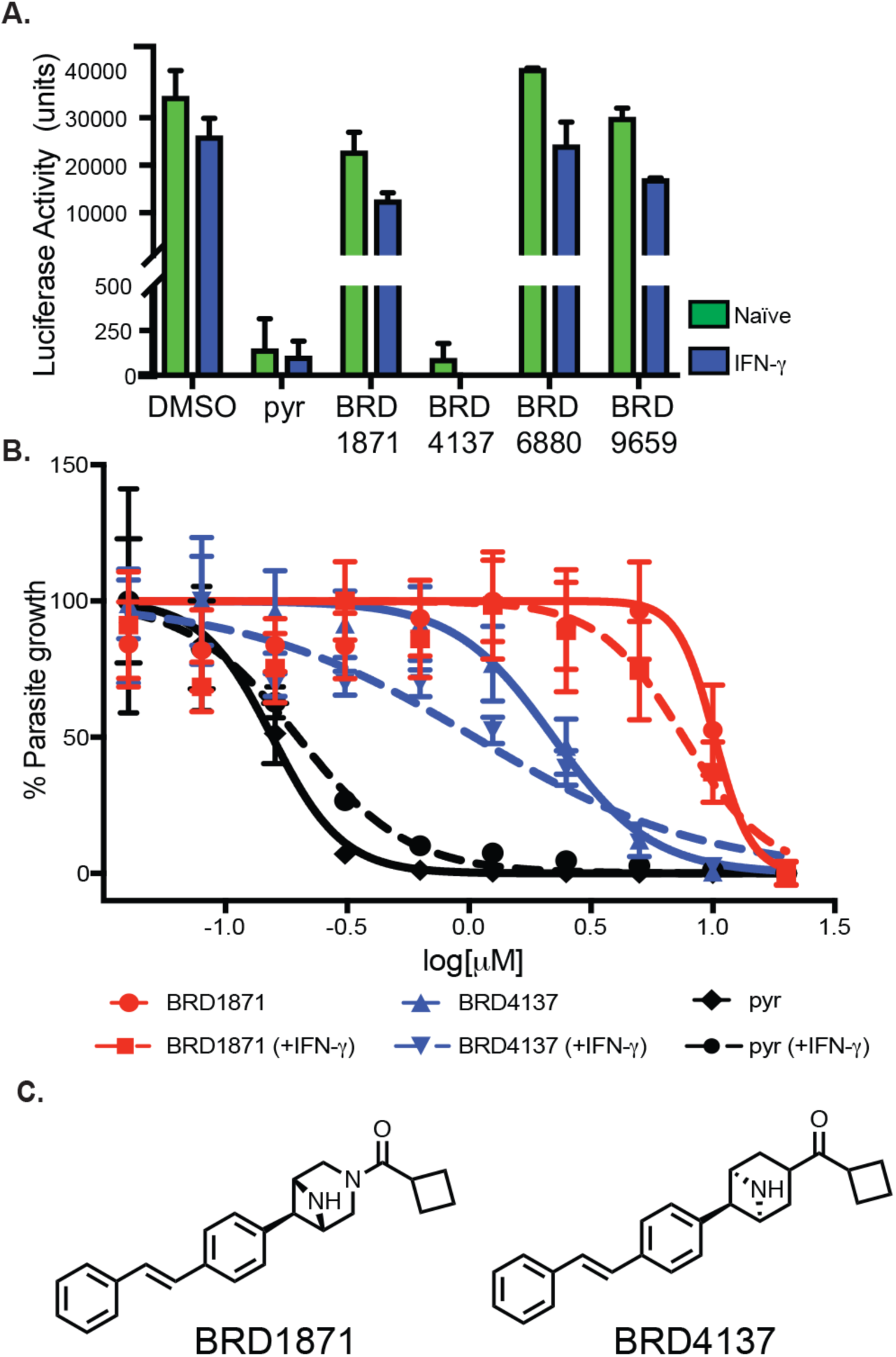
Analysis of BRD1871 and related analogs. **A)** IFN-γ-dependent growth inhibition for parent compound BRD1871 and four analogs within the same chemical series. Green bars = 10 μM compound only (naïve), blue bars = 10 μM compound + 5 U/mL IFN-γ (5U IFN-γ). For comparison, 3 μM pyrimethamine (pyr) was used as a positive control for growth inhibition and DMSO was used as a vehicle control. Average of two independent replicates. **B)** EC_50_ determination generated from 10-point dose response curve in the absence of IFN-γ vs. activated with 5 U/mL IFN-γ (2-fold dilution series with initial dilution at 20 μM). Three biological replicates ± SEM used to generate non-linear regression curve. Complete EC_50_ data are found in Table S3. **C)** The compound structures for bridged bicyclic azetidine compound BRD1871 and a related stereoisomer BRD4137.

Pyrimethamine (black lines) was included to illustrate not all compounds are IFN-γ-dependent (EC_50_ = 0.16 μM naive, solid black line; EC_50_ = 0.20 μM, + 5 U/mL IFN-γ, dashed black line) (Fig. 6B). For compound BRD4137, a single change in the stereochemical properties of BRD1871 resulted in an increased potency against *T. gondii* growth (Fig. 6C), illustrating an important chemical feature for anti-*Toxoplasma* activity of this compound series.

### Analysis of stereoisomers of BRD7522 identifies stereo-selectivity of compound activity

Compound BRD7522 was one of the original hits that showed an EC_50_ that was enhanced 2.1 fold by IFN-γ. BRD7522 contains a bicyclic moiety with a tri-substituted azetidine that has 3 stereocenters (at C1-3) and hence can give rise to a total of 8 possible stereoisomers (Table 1). Most of the stereoisomers showed improved or comparable growth inhibitory activities against *T. gondii* in presence vs. absence of IFN-γ except for BRD6541 and BRD5882 (Table 1). These two compounds have the same configuration (*R*) on the stereocenter at C3 of the azetidine ring while having different configurations at C1 and C2. Changing the C1C2 configuration from *RS*/*SR* to *SS* (BRD7523) yielded similar potency both with and without IFN-γ stimulation (EC_50_ 4.46 μM and 4.14 μM respectively) (Table 1). Switching the C1C2 configuration from *RS*/*SR* to *RR* (BRD0367 and BRD6698) led to slight increases in IFN-γ-dependent growth restriction of the parasite (Table 1). However, the biggest increase in IFN-γ dependent fold changes were observed in the case of BRD4624 and BRD0481 where C2C3 possessed the *SS* configuration (Table 1).

**Table 1.**
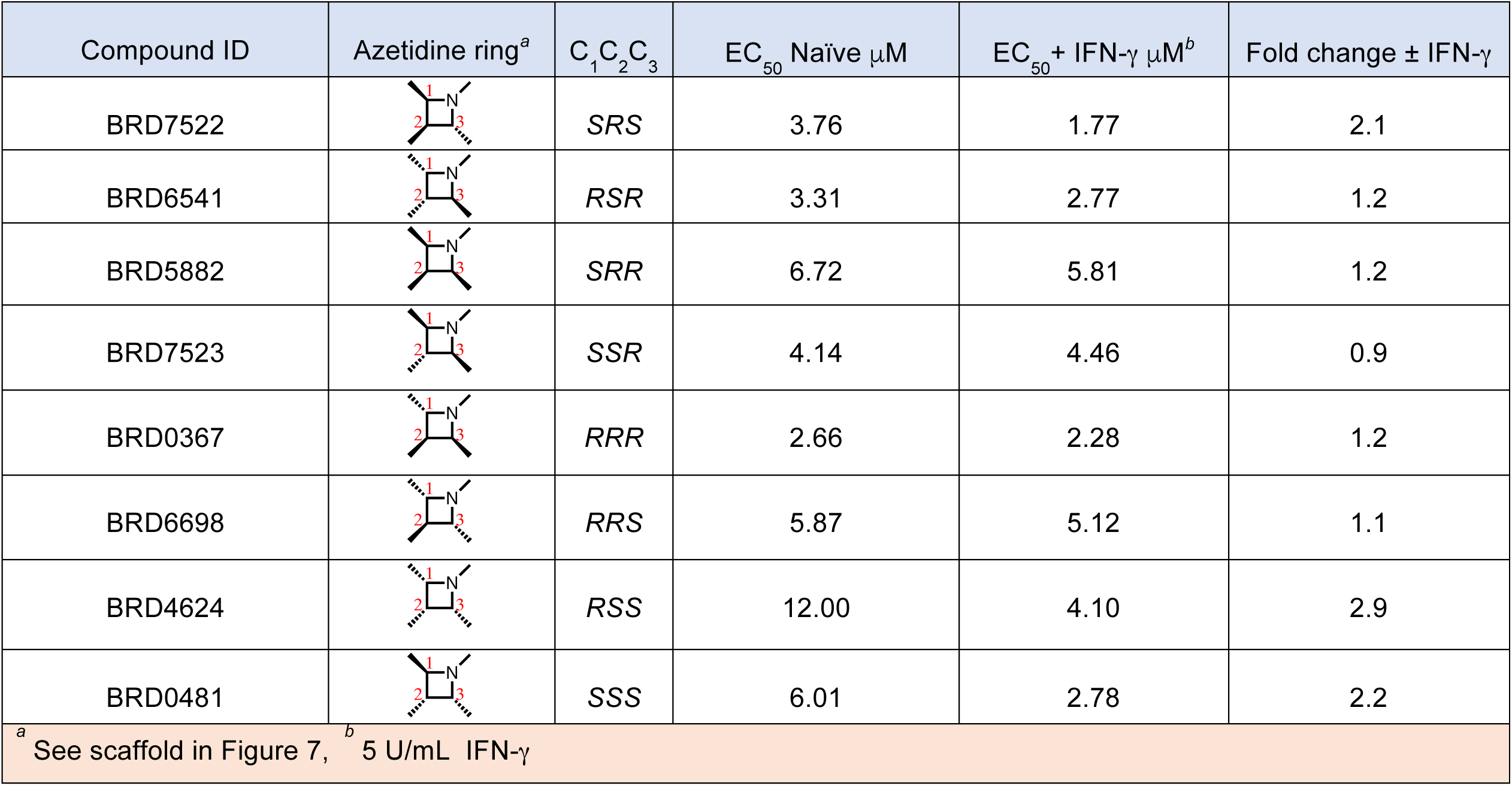
Activities of stereoisomers of BRD7522.

| Compound ID | Azetidine ring<sup>a</sup> | C<sub>1</sub>C<sub>2</sub>C<sub>3</sub> | EC<sub>50</sub> Naïve μM | EC<sub>50</sub> + IFN-γ μM<sup>b</sup> | Fold change ± IFN-γ |
| --- | --- | --- | --- | --- | --- |
| BRD7522 |  | *SRS* | 3.76 | 1.77 | 2.1 |
| BRD6541 |  | *RSR* | 3.31 | 2.77 | 1.2 |
| BRD5882 |  | *SRR* | 6.72 | 5.81 | 1.2 |
| BRD7523 |  | *SSR* | 4.14 | 4.46 | 0.9 |
| BRD0367 |  | *RRR* | 2.66 | 2.28 | 1.2 |
| BRD6698 |  | *RRS* | 5.87 | 5.12 | 1.1 |
| BRD4624 |  | *RSS* | 12.00 | 4.10 | 2.9 |
| BRD0481 |  | *SSS* | 6.01 | 2.78 | 2.2 |
| <sup>a</sup> See scaffold in Figure 7, <sup>b</sup> 5 U/mL IFN-γ |  |  |  |  |  |

A closer look at the stereo-structure activity relationship (SSAR) reveals that the IFN-γ-dependent growth restriction of *T. gondii* is more closely dependent on absolute stereochemistry at position C3 of azetidine ring compared to C1 and C2. Compounds with (*S*) configuration at C3 (e.g. BRD7522, BRD4624, BRD0481, BRD6698) show greater IFN-γ-dependent growth restriction than the corresponding (*RRR*) isomers (e.g. BRD5882, BRD6541, BRD7523, BRD0367) (Fig. 7, Table 1).

**Figure 7:**
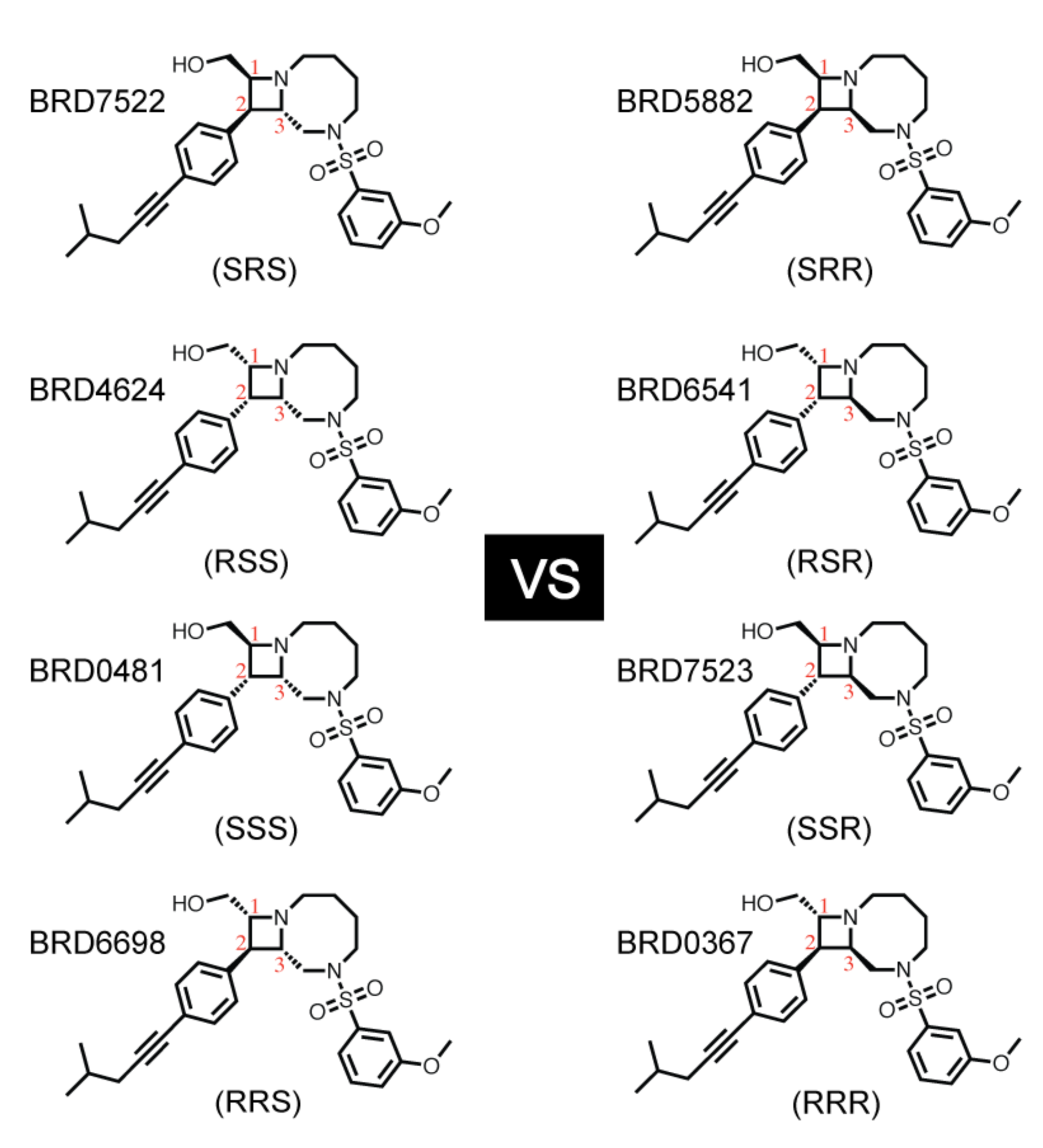
Dependence of *T. gondii* growth inhibition fold change on stereochemical configuration at C3. Compounds with (*SSS*) configuration (left column) exhibit higher dependence on IFN-γ than those with (*RRR*) configuration (right column) as shown in Table 1.

Among the compounds with (*SSS*) configuration, the compound with ‘all-cis’ configuration around the azetidine ring (BRD4624) shows highest IFN-γ-dependent growth restriction (Fig. 7, Table 1).

### Conclusions

We developed a HTS to identify small molecules that exhibit growth inhibition of *T. gondii* that was enhanced by IFN-γ. A number of these inhibitors increased LC3 recruitment to the parasite containing vacuole, consistent with previous reports that a non-canonical autophagy pathway restricts intracellular growth of *T. gondii* (14). However, at least half of the IFN-γ-dependent inhibitors identified here did not work through enhanced LC3 recruitment, as shown by the low correlation between LC3 recruitment and IFN-γ enhancement. Thus in addition to enhancing LC3 recruitment, it appears that IFN-γ can enhance the inhibitory effect of small molecules through other pathways yet to be elucidated. Although the effect of IFN-γ enhancement was modest ranging from 1-3 fold, this property was often consistently seen within a chemical series, suggesting that it is attributable to features of the compound structure. Comparison of one series exemplified by BRD7522 revealed that specific stereoisomers were more strongly associated with inhibition and IFN-γ-dependence, suggesting a specific structural feature of the compounds are responsible for these effects. Hence, synthesis of additional analogs based on the hits discovered here might be exploited to enhance both potency and IFN-γ-dependence. The use of compounds that are synergistic with immune responses may allow more effective treatment at lower drug concentrations, thus minimizing adverse outcomes associated with antibiotic use.

## Experimental Section

### Parasite strains and culture

*Toxoplasma gondii* tachyzoites (ME49-FLUC, type II) (27) were serially passaged in human foreskin fibroblasts (HFF) cultured in complete DMEM medium (DMEM supplemented with 10% fetal bovine serum (FBS), glutamine (10 mM) and gentamycin (10 μg/mL)). For each assay described, infected monolayers were scraped, needle passed (23 ga), and parasites separated from host cell debris using a 3 micron pore polycarbonate filter. HeLa cells were cultured in MEM media that contained 10% FBS, 4 mM L-glutamine and 10 mM HEPES solution. For GFP-LC3 HeLa cell lines, the MEM media described above was supplemented with G418 (50 μg/mL) to maintain the expression plasmid. Cell culture and assays were conducted using a 37°C incubator with 5% CO_2_. Cell cultures were negative for mycoplasma, as determined using the e-Myco plus kit (Intron Biotechnology).

### Luciferase based growth assays: Titration of IFN-γ dependence

HeLa cells were plated in white, clear-bottom 96-well assay plates (Costar #3610) at a density of 5×10^3^ cells/well (200 μL total volume) and allowed to settle overnight. The media was removed and replaced with 100 μL of culture media containing a 2-fold dilution series of recombinant human IFN-γ (R&D Systems, 285-IF-100/CF) (from 100 U/mL to 01. U/ml and including a no IFN-γ control). Tachyzoites of the ME49-FLUC strain were added 24 h post IFN-γ activation at MOI=0.25 (5×10^3^ parasites/well in 100 μL of culture medium, 200 μL total well volume) and allowed to replicate for 72 h post-infection. The luciferase assay was completed according to Luciferase Assay System protocol (Promega, E1501) using the following processing steps: After 72 h, culture medium was dumped and 30 μL of 1x Passive Lysis Buffer (1x PLB, Promega, E1531) was added to each well and incubated for 10 min at room temperature (RT). Plates were read with a Berthold TriStar 941 Multimode Microplate Reader using the following protocol: Inject 100 μL Luciferase Assay Reagent (LAR), shake 1 sec and read luminescence at 10 sec post injection.

### Scaling the assay to 384 well plate format

A high throughput screening (HTS) assay was miniaturized from a 96-well to a 384-well plate format to allow for more efficient screening. On day 1, HeLa cells (ATCC CCL-2) were plated in 384-well white plates at a density of 1,000 cells/well (20 μL total volume) in DMEM with 10% Fetal bovine serum (Hyclone) and allowed to settle. The following day, cells were activated with IFN-γ (1.25 U/mL, 10 μL volume) and returned to the incubator for 24 h. On day 3, 100 nL of library compound was added to each well of the assay plates using a pin transfer method yielding a concentration of 20 μM compound in each assay well. Freshly egressed ME49-FLUC parasites were diluted to 4,000 parasites/well (MOI=1, 20 μL volume) and added to each well, resulting in a 10 μM final compound concentration (50 μL, final well volume). For compound transfer/infection runs greater than 2 h, fresh parasites were harvested every 2 h and resupplied to the automation platform to ensure consistent infection. On day 6, 30 μL of Steady-Glo reagent (Promega) was added to each well and after a brief incubation the Luminescence was detected on an Envision multi-mode plate reader (Perkin Elmer). To test for possible toxicity, HeLa cells were incubated with 20 μM compound (50 μL, final well volume) for 72 h and viability was assessed by measuring steady state ATP levels using the CellTiter-Glo reagent (Promega). Compounds that showed a reduction in ATP levels by 50% were eliminated.

### Compound libraries

Compounds were chosen from a diverse unbiased collection of compounds drawn from the commercial libraries, including WuXi, Asinex, ChemBridge, ChemDiv, Enamine, MayBridge, Sigma, the Broad Institute Diversity Oriented Synthesis (DOS) library and a FDA repurposing collection of FDA approved and clinical candidates assembled at the Broad Institute (28). All compounds were assembled and sourced at the Broad Institute. At the time of purchase, these commercial collections were prescreened using a computation chemistry algorithm to identify and exclude all PAIN compounds. A computational chemistry search was used to identify a subset of plates within the larger composite chemical library to maximize the chemical diversity of compounds screened in this HTS effort. At the time of pulling the compounds out of storage for retesting at dose, the compounds were put through a quality control step to ensure all compounds were of the correct mass and greater than 85% pure.

### Luciferase based growth assays: validation of hits

*Single point assays.* White, clear-bottom 96-well assay plates were prepared as described above, with the following modifications. HeLa cells were activated for 24 h with low dose IFN-γ (5 U/mL,100 μL/well) or naïve (unactivated, 100 μL/well). At 24 h after addition of IFN-γ, 50 μL of 4x compound were added to each well and incubated for 4 h prior to the addition of ME49-FLUC parasites (5×10^3^ parasites/well in 50 μL, 200 μL final well volume). Compounds were provided as 10 mM stock in 100% DMSO, diluted to 4x concentration in culture media (0.4% DMSO) and added to each assay well to reach 5 (0.05% DMSO final) or 10 μM (0.1% DMSO final) final compound concentration. Luciferase assays were completed at 72 h post infection using the following parameters: Medium was removed and replaced with 40 μL of 1x PLB/well, incubated for 5 min at RT and read on the Cytation 3 Imaging Multi-Mode Imager using these parameters: dispense 100 μL LAR, shake 1 sec, read well at 10 sec post injection.

*Multi-point dose response (EC_50_ determination).* White, clear bottom 96-well assay plates were set up as described above with the following modifications. Only the inner 60 wells were used for the assay in order to reduce variability that results from evaporation during incubation. Dilution of compound stock is as described above, with all wells containing a final concentration of 0.1% DMSO. Compounds were compared using 2-fold-serial dilution from 20 μM to 0.04 μM in the absence vs. presence of IFN-γ (5 or 10 U/ml). DMSO (vehicle control) and pyrimethamine (2.5 μM, positive control) were included in the outside wells of all plates as controls. Liquid handling (serial dilutions, compound dilutions, cell feeding, plate-to-plate transfers, etc) utilized a Dual Pod Biomek FX. Luciferase assays were performed using the integrated and automated platform (Beckman Coulter). The SAMI EX software was used to design and execute the screening assay and enabled efficient and uniform assay execution across all the assay plates. All steps and assays were completed at the High Throughput Screening Center at Washington University School of Medicine as described above.

### Image based GFP-LC3 recruitment assay

GFP-LC3 HeLa cells were plated in 96-well, μClear black microplates (Greiner Bio-One, #655090) at a density of 1.8×10^4^ cells/well and incubated overnight at 37°C. Cells were activated for 24 h with low dose IFN-γ (5 U/mL, 100 μL/well) or left naïve (100 μL/well), followed by a 4 h pretreatment with indicated compounds (5 μM, final concentration, see above for stock compound dilution and final DMSO % in each well) prior to the addition of ME49-FLUC parasites (5×10^4^ parasites/well in 50 μL; 200 μL/well final volume). After 2 h, wells were rinsed 3 times in culture media to remove extracellular parasites and returned to normal culture media. Cells were fixed 6 h post-infection with 4% formaldehyde, permeabilized with 0.05% saponin and stained for indirect immunofluorescence. GFP-LC3 was localized using a mouse monoclonal antibody 3E6 (Life Technologies) and detected by Alexa Fluor 488 (Invitrogen). Parasite containing vacuoles were localized using rabbit polyclonal serum against type I parasites (RH) and detected using Alexa Fluor 594 (Invitrogen). Images were collected on a Cytation 3 imager and positive recruitment events were defined as overlap of GFP-LC3 signal (green) with *T. gondii* (red) signal that met minimum thresholds for a positive interaction (green: ≥55,000 units; red; size requirement range from 5 to 11 microns and signal >5,000 units).

### Statistics

All results are presented as the average of two or more biological replicates. Linear regression analysis and dose-response inhibition (Log(inhibitor) vs. normalized response – variable slope) or (Log(inhibitor) vs normalized response) were performed in Prism 7 (GraphPad Software, Inc.).

### Ancillary Information

#### Supporting Information

*Corresponding author information:* Jose R. Perez, L. David Sibley

*Data access*: Screening data have been submitted to PubChem and the AID reference number is pending.

## Acknowledgement

We thank Dr. Anne Paredes, Department of Pathology and Immunology, Washington University, for coordinating interactions among the research groups and Dr. Maxine Ilagan, High-Throughput Screening Center at Washington University School of Medicine, for assistance with compound screening. Supported by the National Institute of Allergy and Infectious Diseases of the National Institutes of Health under Award Number U19AI109725.

## Author contributions

JBR, KLC, JRP, designed or performed screening assays; JBR, KLC, SS analyzed primary screening data; CM, JB, RH, SS, SLS, provided advice on compound selection or activity; JBR, KLC, JRP, generated figures; SLS, HWV, JRP, LDS provided scientific advice and supervision; JBR, JRP, KLC, LDS wrote the manuscript with input from all authors.

## Abbreviations used

*IFN-γ*: interferon gamma
*PYR*: pyrimethamine
*LC3*: light chain 3
*FLUC*: firefly luciferase
*EC*: effective concentration
*GFP*: green fluorescent protein

## References

1. Dubey JP. 2010. Toxoplasmosis of animals and humans. CRC Press, Boca Raton.

2. Knoll LJ, Tomita T, Weiss LM. 2014. Bradyzoite development, p. 521-551. In Weiss LM, Kim K (ed.), Toxoplasma gondii: The model apicomplexan: perspectives and methods, 2nd ed. Academic Press, New York.

3. Hide G. 2016. Role of vertical transmission of Toxoplasma gondii in prevalence of infection. Expert review of anti-infective therapy 14:335–344.

4. Su C, Evans D, Cole RH, Kissinger JC, Ajioka JW, Sibley LD. 2003. Recent expansion of Toxoplasma through enhanced oral transmission. Science 299:414–416.

5. Jones JL, Dubey JP. 2010. Waterborne toxoplasmosis--recent developments. Exp Parasitol 124:10–25.

6. Jones JL, Dubey JP. 2012. Foodborne Toxoplasmosis. Clin Infect Dis 55:864–851.

7. Torgerson PR, Mastroiacovo P. 2013. The global burden of congenital toxoplasmosis: a systematic review. Bull World Health Organ 91:501–508.

8. Pappas G, Roussos N, Falagas ME. 2009. Toxoplasmosis snapshots: global status of *Toxoplasma gondii* seroprevalence and implications for pregnancy and congenital toxoplasmosis. Int J Parasitol 39:1385–1394.

9. Montoya JG, Liesenfeld O. 2004. Toxoplasmosis. Lancet 363:1965–1976.

10. MacMicking JD. 2012. Interferon-inducible effector mechanisms in cell-autonomous immunity. Nat Rev Immunol 12:367–382.

11. Suzuki Y, Orellana MA, Schreiber RD, Remington JS. 1988. Interferon-g: the major mediator of resistance against *Toxoplasma gondii*. Science 240:516–518.

12. Yap GS, Sher A. 1999. Effector cells of both nonhemopoietic and hemopoietic origin are required for interferon (IFN)-gamma- and tumor necrosis factor (TNF)-alpha-dependent host resistance to the intracellular pathogen, *Toxoplasma gondii*. J. Exp. Med. 189:1083–1091.

13. Saeij JP, Frickel EM. 2017. Exposing Toxoplasma gondii hiding inside the vacuole: a role for GBPs, autophagy and host cell death. Curr Opin Microbiol 40:72–80.

14. Selleck EM, Orchard RC, Lassen KG, Beatty WL, Xavier RJ, Levine B, Virgin HW, Sibley LD. 2015. A Noncanonical Autophagy Pathway Restricts Toxoplasma gondii Growth in a Strain-Specific Manner in IFN-gamma-Activated Human Cells. MBio 6:e01157–01115.

15. Pfefferkorn ER. 1984. Interferon-gamma blocks the growth of *Toxoplasma gondii* in human fibroblasts by inducing the host to degrade tryptophan. Proc. Natl. Acad. Sci. (USA) 81:908–912.

16. Haldar AK, Foltz C, Finethy R, Piro AS, Feeley EM, Pilla-Moffett DM, Komatsu M, Frickel EM, Coers J. 2015. Ubiquitin systems mark pathogen-containing vacuoles as targets for host defense by guanylate binding proteins. Proc Natl Acad Sci U S A 112:E5628–5637.

17. Clough B, Wright JD, Pereira PM, Hirst EM, Johnston AC, Henriques R, Frickel EM. 2016. K63-Linked Ubiquitination Targets Toxoplasma gondii for Endo-lysosomal Destruction in IFNgamma-Stimulated Human Cells. PLoS Pathog 12:e1006027.

18. Niedelman W, Sprokholt JK, Clough B, Frickel EM, Saeij JP. 2013. Cell death of gamma interferon-stimulated human fibroblasts upon *Toxoplasma gondii* infection induces early parasite egress and limits parasite replication. Infect Immun 81:4341–4349.

19. Jones LA, Alexander J, Roberts CW. 2006. Ocular toxoplasmosis: in the storm of the eye. Parasite Immunol. 28:635–642.

20. Wei HX, Wei SS, Lindsay DS, Peng HJ. 2015. A Systematic Review and Meta-Analysis of the Efficacy of Anti-Toxoplasma gondii Medicines in Humans. PLoS One 10:e0138204.

21. McCabe RE. 2001. Antitoxoplasma chemotherapy, p. 319-359. *In* Joynson DHM, Wreghitt TG (ed.), Toxoplasmosis: a comprehensive clinical guide. Cambridge Univ. Press, Cambridge.

22. Araujo CH, Remington J. 1991. Synergistic activity of azithromycin and gamma interferon in murine toxoplasmosis. Antimicrob. Agents Chem. 335:1672–1673.

23. Dupont CD, Christian DA, Hunter CA. 2012. Immune response and immunopathology during toxoplasmosis. Semin Immunopathol 34:793–813.

24. Schreiber SL, Nicolaou KC, Davies K. 2002. Diversity-oriented organic synthesis and proteomics. New frontiers for chemistry & biology. Chem Biol 9:1–2.

25. Schreiber SL. 2000. Target-oriented and diversity-oriented organic synthesis in drug discovery. Science 287:1964–1969.

26. Xia X, Owen MS, Lee RE, Gaudet S. 2014. Cell-to-cell variability in cell death: can systems biology help us make sense of it all? Cell Death Dis 5:e1261.

27. Tobin CM, Knoll LJ. 2012. A patatin-like protein protects Toxoplasma gondii from degradation in a nitric oxide-dependent manner. Infect Immun 80:55–61.

28. Corsello SM, Bittker JA, Liu Z, Gould J, McCarren P, Hirschman JE, Johnston SE, Vrcic A, Wong B, Khan M, Asiedu J, Narayan R, Mader CC, Subramanian A, Golub TR. 2017. The Drug Repurposing Hub: a next-generation drug library and information resource. Nat Med 23:405–408.

